# Human knockouts in a cohort with a high rate of consanguinity

**DOI:** 10.1101/031518

**Authors:** Danesh Saleheen, Pradeep Natarajan, Wei Zhao, Asif Rasheed, Sumeet Khetarpal, Hong-Hee Won, Konrad J. Karczewski, Anne H O’Donnell-Luria, Kaitlin E. Samocha, Namrata Gupta, Mozzam Zaidi, Maria Samuel, Atif Imran, Shahid Abbas, Faisal Majeed, Madiha Ishaq, Saba Akhtar, Kevin Trindade, Megan Mucksavage, Nadeem Qamar, Khan Shah Zaman, Zia Yaqoob, Tahir Saghir, Syed Nadeem Hasan Rizvi, Anis Memon, Nadeem Hayyat Mallick, Mohammad Ishaq, Syed Zahed Rasheed, Fazal-ur-Rehman Memon, Khalid Mahmood, Naveeduddin Ahmed, Ron Do, Daniel G. MacArthur, Stacey Gabriel, Eric S. Lander, Mark J. Daly, Philippe Frossard, John Danesh, Daniel J. Rader, Sekar Kathiresan

## Abstract

A major goal of biomedicine is to understand the function of every gene in the human genome.^1^ Null mutations can disrupt both copies of a given gene in humans and phenotypic analysis of such ‘human knockouts’ can provide insight into gene function. To date, comprehensive analysis of genes knocked out in humans has been limited by the fact that null mutations are infrequent in the general population and so, observing an individual homozygous null for a given gene is exceedingly rare.^2,3^ However, consanguineous unions are more likely to result in offspring who carry homozygous null mutations. In Pakistan, consanguinity rates are notably high.^4^ Here, we sequenced the protein-coding regions of 7,078 adult participants living in Pakistan and performed phenotypic analysis to identify homozygous null individuals and to understand consequences of complete gene disruption in humans. We enumerated 36,850 rare (<1 % minor allele frequency) null mutations. These homozygous null mutations led to complete inactivation of 961 genes in at least one participant. Homozygosity for null mutations at *APOC3* was associated with absent plasma apolipoprotein C-III levels; at *PLAG27,* with absent enzymatic activity of soluble lipoprotein-associated phospholipase A2; at *CYP2F1,* with higher plasma interleukin-8 concentrations; and at either *A3GALT2* or *NRG4,* with markedly reduced plasma insulin C-peptide concentrations. After physiologic challenge with oral fat, *APOC3* knockouts displayed marked blunting of the usual post-prandial rise in plasma triglycerides compared to wild-type family members. These observations provide a roadmap to understand the consequences of complete disruption of a large fraction of genes in the human genome.

## Main Text

We studied adult participants in the Pakistan Risk of Myocardial Infarction Study (PROMIS) designed to understand the determinants of cardiometabolic diseases in South Asians.^5^ Consanguineous marriages have been common in this region of South Asia for many generations.^6^ In PROMIS, 38.3% of participants reported that their parents were cousins and 38.1% reported themselves being married to a cousin. An expectation from consanguinity is long regions of autozygosity, defined as homozygous loci identical by descent.^7^ Using genome-wide genotyping data available in 17,744 PROMIS participants, we quantified the length of runs of homozygosity, defined as homozygous segments at least 1.5 megabases long. We compared the lengths of runs of homozygosity among PROMIS participants with that seen in other populations from the International HapMap3 project. Median length of genome-wide homozygosity among PROMIS participants was 6–7 times higher than participants of European (CEU, TSI) (P = 3.6 × 10”^37^), East Asian (CHB, JPT, CHD) (P = 5.4 × 10^−48^) and African ancestries (YRI, MKK) (P = 1.3 × 10^−40^), respectively (**Fig. 1**).

**Fig 1.**
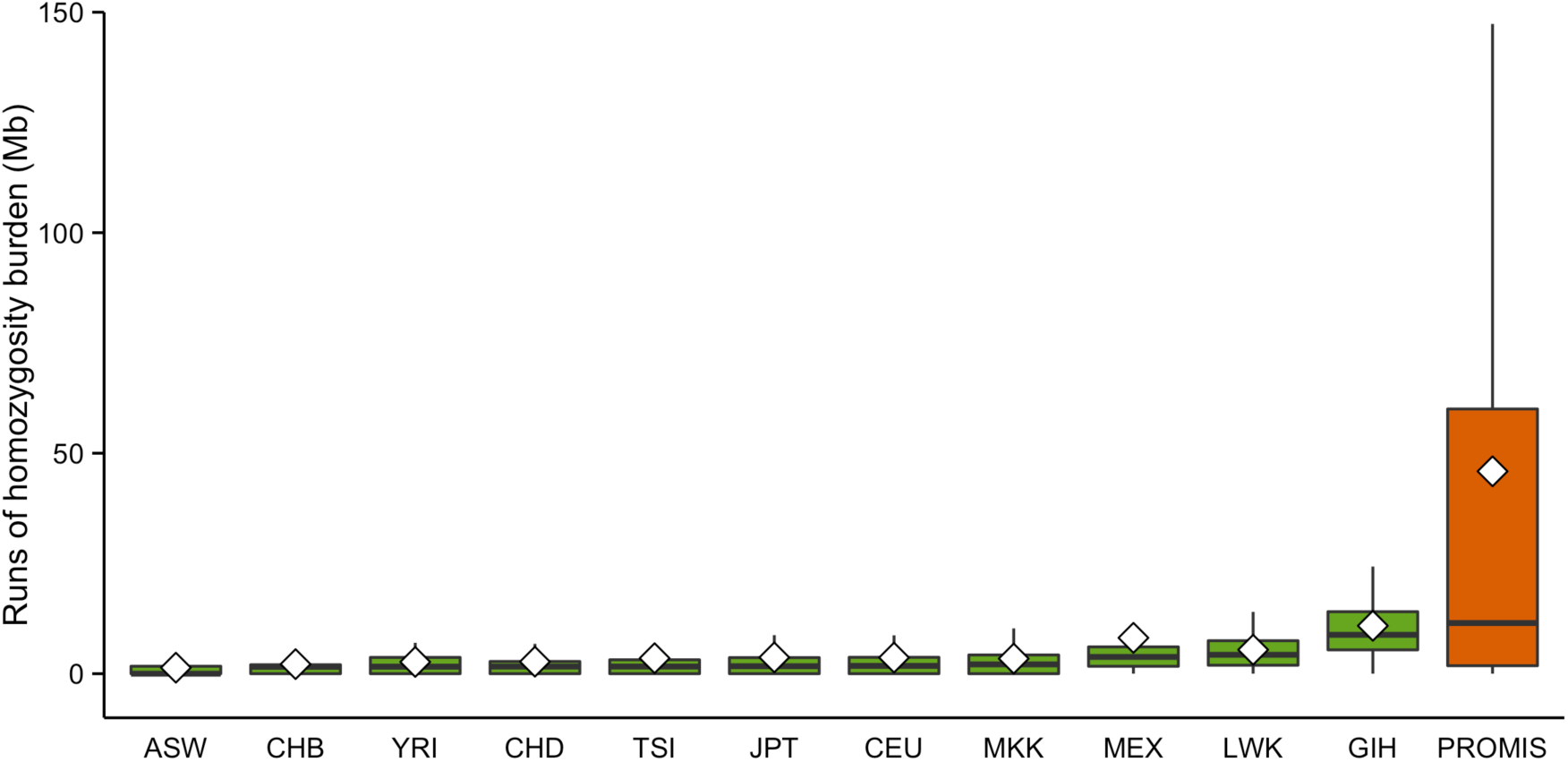
PROMIS participants have an excess burden of runs of homozygosity compared with other populations.

In order to characterize the burden of rare homozygous null alleles, we performed whole exome sequencing in 7,078 PROMIS participants (**Table 1**). Across all participants, 1,303,689 protein-coding and splice-site sequence variants in 20,008 autosomal genes passed variant quality control metrics. Of these, 47,656 mutations across 13,645 autosomal genes were annotated as null (nonsense, frameshift, or canonical splice-site). To increase the probability that mutations annotated as null are *bona fide*, we removed nonsense and frameshift mutations occurring within the last 5% of the transcript and within exons flanked by non-canonical splice sites, splice site mutations at small (<15 bp) introns, at non-canonical splice sites, and where the purported null allele is observed across primates. Common null alleles are less likely to exert strong functional effects as they are less constrained by purifying selection; thus, we limit our analyses in the rest of the manuscript to null mutations with a minor allele frequency (MAF) of < 1%.

**Table 1.** Baseline characteristics of exome sequenced study participants.

| **Characteristic** | **Value** |
| --- | --- |
|  | **(n = 7,078)** |
| **Age (yrs) – mean (sd)** | 50.7 (9.1) |
| **Women – no. (%)** | 1,249 (17.6 %) |
| **Parents closely related – no. (%)** | 2,710 (38.3 %) |
| **Spouse closely related – no. (%)** | 2,697 (38.1 %) |
| **Ethnicity – no. (%)** |  |
| **Urdu** | 2,647 (37.4 %) |
| **Punjabi** | 2,332 (32.9 %) |
| **Sindhi** | 797 (11.3 %) |
| **Pathan** | 416 (5.9 %) |
| **Memon** | 108 (1.5 %) |
| **Gujrati** | 89 (1.3 %) |
| **Balochi** | 85 (1.2 %) |
| **Other** | 604 (8.5 %) |
| **Hypertension – no. (%)<sup>\*</sup>** | 3,936 (55.6 %) |
| **Hypercholesterolemia – no. (%)<sup>†</sup>** | 1,857 (26.2 %) |
| **Diabetes mellitus – no. (%)<sup>‡</sup>** | 2,904 (41.0 %) |
| **Coronary heart disease – no. (%)<sup>§</sup>** | 3,046 (43.0 %) |
| **Smoking – no. (%)** | 2,814 (39.8 %) |
| **BMI (m/kg<sup>2</sup>) – mean (sd)** | **25.9 (4.3)** |

\*Hypertension defined as systolic blood pressure $\geq 140$ mmHg, diastolic blood pressure $\geq 90$ mmHg, or antihypertensive treatment.
†Hypercholesterolemia defined as serum total cholesterol $> 240$ mg/dL, lipid lowering therapy or self-report.
‡Diabetes defined as fasting blood glucose $\geq 126$ mg/dL, or HbA1c $> 6.5$ %, oral hypoglycemics, insulin treatment, or self-report.
§Coronary heart disease defined as history of myocardial infarction as determined by clinical symptoms with typical EKG findings or elevated serum troponin I.
||Smoking defined as active current or prior tobacco smoking.

Applying these criteria, we generated a set of 36,850 null mutations across 12,131 autosomal genes.^8^ The site-frequency spectrum for these null mutations revealed that the majority was seen only in one or a few individuals (**Fig. 2**).

**Fig 2.**
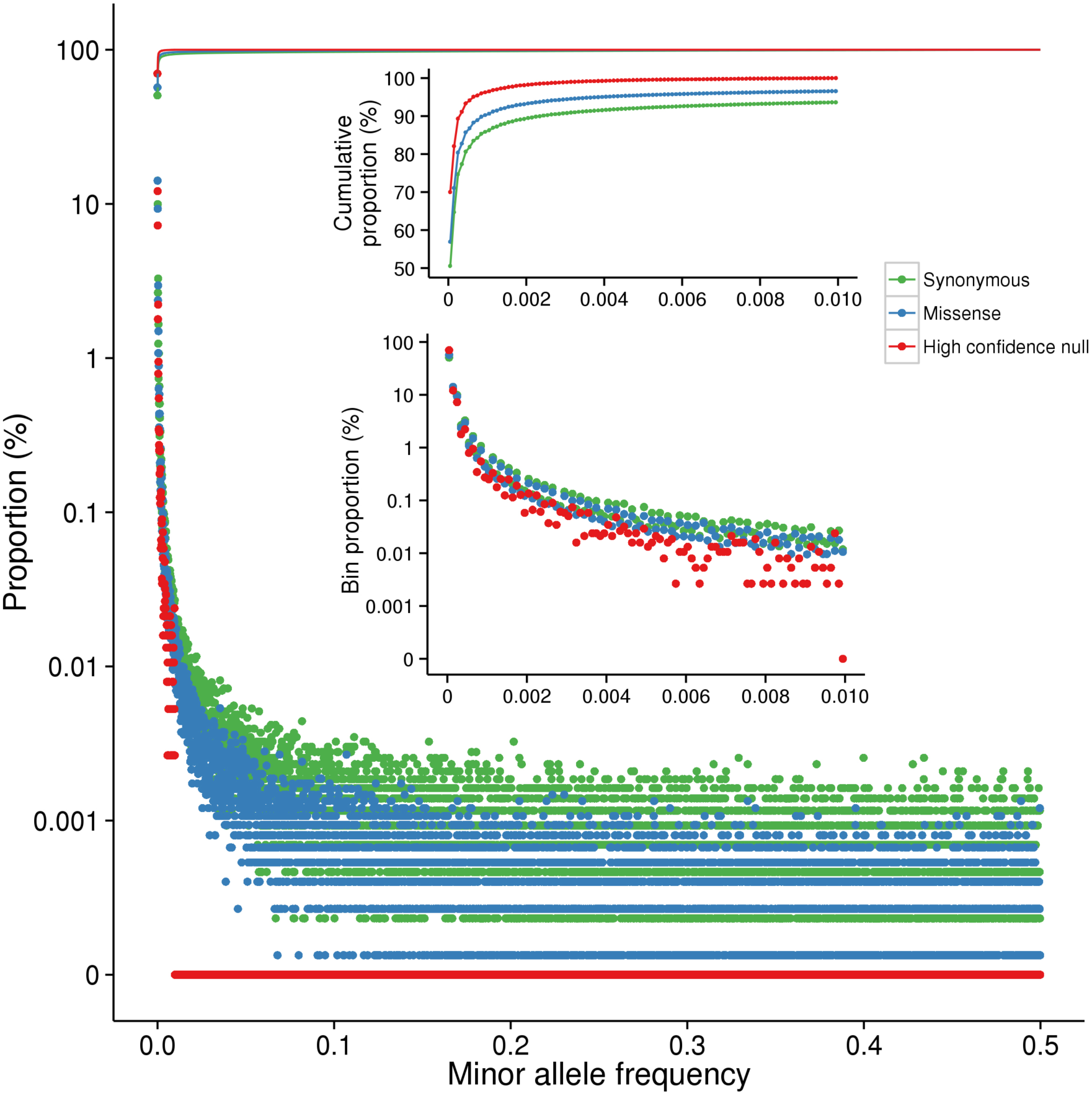
Null mutations are typically seen in very few individuals.

We compared the coefficient of inbreeding (F coefficient) in PROMIS participants with that of 15,249 individuals from outbred populations of European or African American ancestry. The F coefficient estimates the excess homozygosity compared with an estimated outbred ancestor. PROMIS participants had a 4-fold higher median inbreeding coefficient compared to outbred populations (0.016 v 0.0041; *P* < 2 × 10^−16^) (**Fig. 3A**). Additionally, those in PROMIS who reported that their parents were closely related had even higher median inbreeding coefficients than those who did not (0.024 v 0.013; *P* < 2 × 10^−16^).

**Fig. 3.**
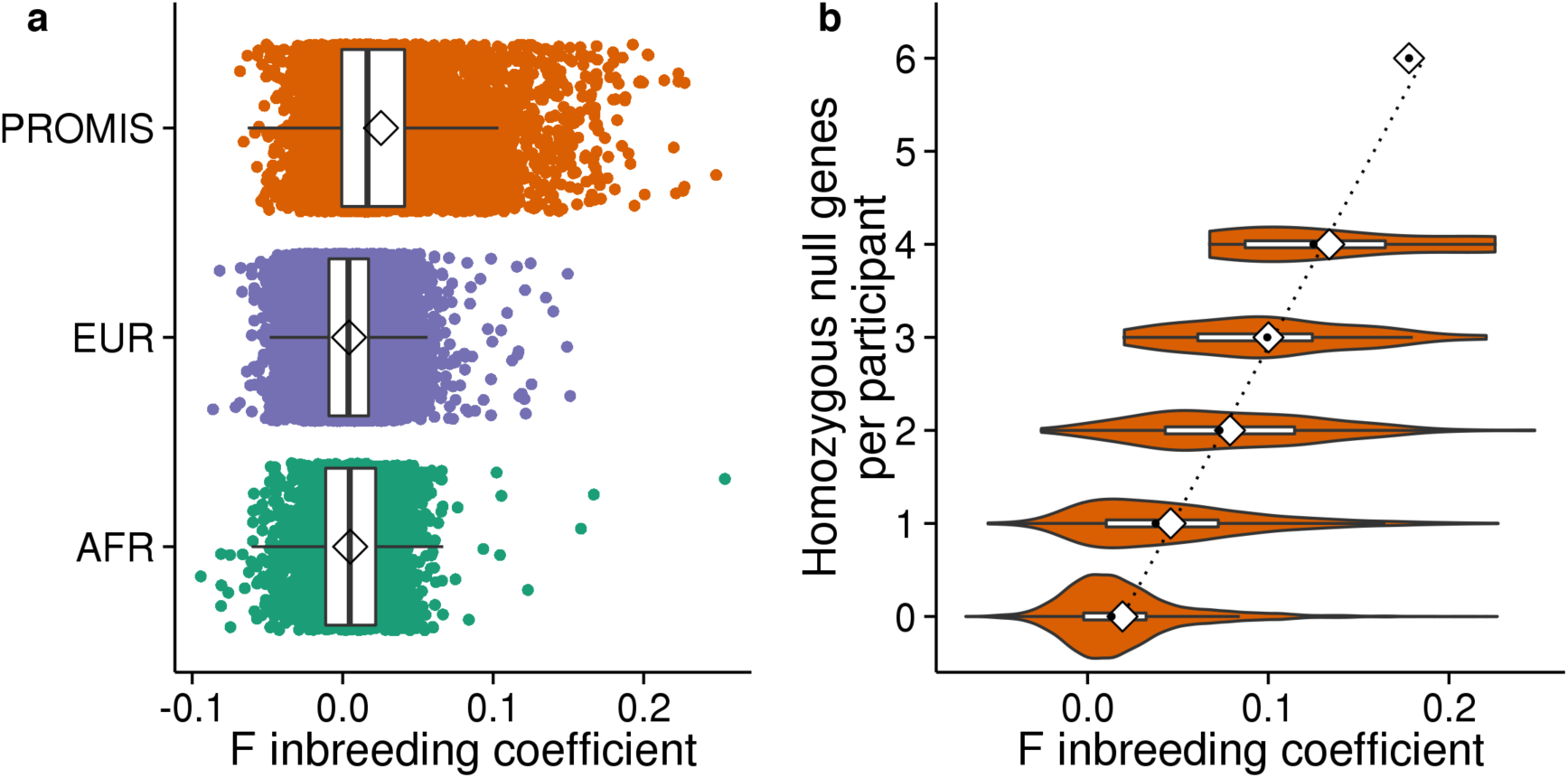
Homozygous null burden in PROMIS is driven by excess autozygosity.

Across all 7,078 PROMIS participants, both copies of 961 distinct genes were disrupted due to null mutations with MAF < 1%. A full listing of all 961 genes knocked out, the number of knockout participants for each gene, and the specific null mutation(s) are provided in **Supplementary Table 1**. 697 (72.5 %) of the genes were knocked out only in one participant (**Supplementary Figure 1**). About 1 in 5 sequenced participants (1,306 individuals, 18.4 %) had at least one gene knocked by a homozygous null mutation with MAF < 1%. 1,081 of these 1,306 individuals (82.8 %) had complete deficiency for one gene, but a minority of participants were knockouts for more than one gene and one participant had six genes completely inactivated. The F inbreeding coefficient was correlated with the number of homozygous null genes present in each individual. (Spearman r = 0.29; *P* = 3 × 10^−133^) (**Fig. 3B**).

We tested the hypothesis that genes observed in the homozygous null state in PROMIS participants are under less evolutionary constraint. We calculated the probability of being loss-of-function intolerant (at >90% threshold) for each gene (see Methods) ^9,10^ and. compared this to 961 randomly selected genes. The observed 961 homozygous null genes were less likely to be classified as highly constrained (odds ratio 0.10; 95% CI 0.095, 0.11; P < 1 × 10^−10^).

We next sought to understand the phenotypic consequences of complete disruption of any of these 961 genes. We applied two approaches. First, for 264 genes where two or more participants were homozygous null, we conducted an association screen against a panel of 201 phenotypic traits (**Supplementary Table 2**). Second, at a single gene, we recalled participants based on genotype across three classes (‘wild-type’, heterozygous null, and homozygous null) and performed provocative physiologic testing.

At 264 genes where two or more participants were homozygous null, we performed association analyses to determine whether homozygous null mutation status was associated with variation in any of 201 traits. For quantitative traits, we compared mean trait values in homozygous null carriers with non-carriers. For dichotomous traits, we performed logistic regression with trait status as the outcome variable and homozygous null carrier status as the predictor variable. Details of covariate adjustments are presented in the Methods. Across quantitative and dichotomous traits, this resulted in the analysis of 15,263 gene-trait pairs and thus, we set Bonferroni-adjusted significance threshold at *P* = 3 × 10^−6^.

The quantile-quantile plot of expected versus observed association results shows an excess of highly significant results without systematic inflation (**Supplementary Figure 2**). Association results surpassed the Bonferroni significance threshold for 14 gene-trait pairs (**Supplementary Table 3**). Below, we highlight four associations demonstrating examples of 1) confirmed biochemical deficiency with two independent assays (*PLA2G7*), 2) gene-biomarker association (*CYP2F1*), 3) pure recessive model of association (*A3GALT2*), and 4) confirmed biochemical deficiency with gene-biomarker association (*APOC3*).

Lipoprotein-associated phospholipase A2 (Lp-PLA2, encoded by *PLA2G7*) hydrolyzes oxidatively-modified polyunsaturated fatty acids producing lysophosphatidylcholine and oxidized nonesterified fatty acids. Higher soluble Lp-PLA2 activity has been correlated with higher risk for cardiovascular disease.^11^ At *PLA2G7,* we identified two participants homozygous for *PLA2G7* c.663+1G>A. When compared with non-carriers, c.663+1G>A homozygotes have markedly lower Lp-PLA2 enzyme as well as activity (−266 ng/ml, *P* = 7 × 10^−5^ for mass; −245 nmol/ml/min, *P* = 2 × 10^−7^ for activity) whereas the 102 heterozygotes had an intermediate effect (−107 ng/ml, *P* = 4 × 10^−26^ for mass; −115 nmol/ml/min, *P* = 3 × 10^−58^ for activity) (**Supplementary Figure 3a-b**).

Cytochrome P450 2F1 (encoded by *CYP2F1*) is primarily expressed in the lung and metabolizes pulmonary-selective toxins, such as cigarette smoke, and thus, modulates the expression of environmentally-associated pulmonary diseases.^12^ At *CYP2F1,* we identified two participants homozygous for a splice-site mutation, c.1295–2A>G. When compared with non-carriers, c.1295–2A>G homozygotes displayed higher soluble interleukin 8 concentrations (+40.3 %; *P* = 3 × 10^−6^) (**Supplementary Figure 4**). *CYP2F1* c.1295–2A>G heterozygotes (n = 3 assayed for interleukin 8) had a more modest effect (+10.7 %; *P* = 2 × 10^−4^). Interleukin 8 is a mediator of acute pulmonary inflammation.^13^

Alpha-1,3-galactosyltransferase 2 (encoded by *A3GALT2*) catalyzes the formation of the Gal-*α*1–3Galβ1–4GlcNAc-R (*α*-gal) epitope; the biological role of this enzyme in humans is uncertain.^14^ At *A3GALT2,* we identified two participants homozygous for a frameshift mutation, p.Thr106SerfsTer4. Compared with non-carriers, p.Thr106SerfsTer4 homozygotes had both reduced fasting insulin C-peptide (−97.4%; *P* = 6 × 10^−12^) as well as total fasting insulin concentrations (−91.9%; *P* = 1 × 10^−4^). Such an association was only observed in the homozygous state (**Supplementary Figure 5**). *A3galt^-/-^* mice and pigs have recently been shown to have glucose intolerance.^15,16^

To understand if the identification of only a single homozygote in PROMIS may still be informative, we performed a complementary analysis, focusing on those with the most extreme standard Z scores (|Z score| > 5) and requiring that there be evidence for association in heterozygotes as well (see Methods). This procedure highlighted neureglin 4 (NRG4), a member of the epidermal growth factor family extracellular ligands which is highly expressed in brown fat, particularly during adipocyte differentiation.^17,18^ At *NRG4,* we identified a single participant homozygous for a frameshift mutation, p.Ile75AsnfsTer23, who had nearly absent fasting insulin C-peptide concentrations (−99.3 %; *P* = 7 × 10^−11^). When compared with non-carriers, heterozygotes for *NRG4* p.Ile75AsnfsTer23 (n = 7) displayed 54.5 % reduction in insulin C-peptide (*P* = 6 × 10^−3^). Mice homozygous deleted for *Nrg4* have recently been shown to have glucose intolerance.^18^

Apolipoprotein C-III (apoC-III, encoded by *APOC3*) is a major protein component of chylomicrons, very low-density lipoprotein cholesterol, and high-density lipoprotein cholesterol.^19^ We and others recently reported that null mutations in heterozygous form lower plasma triglycerides as well as risk for coronary heart disease.^20,21^ In both published studies, no *APOC3* homozygotes were identified despite study of nearly 200,000 participants from the U.S. and Europe. However, in this study of about 7,000 Pakistanis, we identified four participants homozygous for *APOC3* p.Arg19Ter. When compared with non-carriers, p.Arg19Ter homozygotes displayed near-absent plasma apoC-III protein (−89.4 %, *P* = 5 × 10^−24^), lower plasma triglyceride concentrations (−61.7 %, *P* = 4 × 10^−4^), higher high-density lipoprotein (HDL) cholesterol (+28.1 mg/dL, *P* = 6 × 10^−9^); and similar levels of low-density lipoprotein (LDL) cholesterol (*P* = 0.11) (**Fig. 4a-d**).

**Fig 4.**
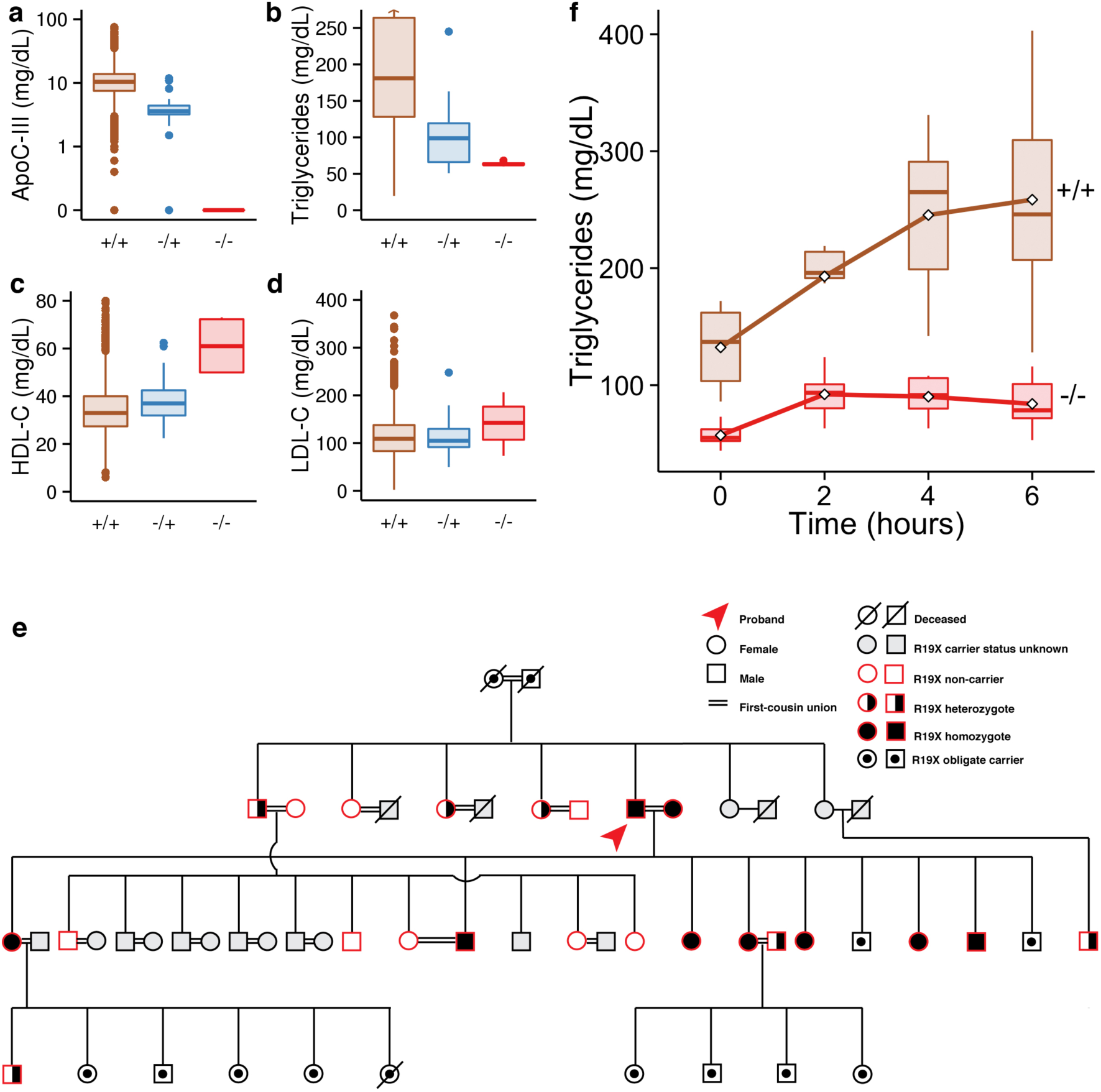
*APOC3* null homozygotes have diminished fasting triglycerides and blunted post-prandial lipemia.

ApoC-III functions as a brake on the metabolism of dietary fat and thus, the complete lack of this protein should promote handling of ingested fat. The availability of humans completely deficient in APOC3 allowed us to test this hypothesis directly. We re-contacted one homozygous null proband, his wife, and 27 of his first-degree relatives for genotyping and physiologic investigation. Surprisingly, we found that the proband’s wife was also a null homozygote, leading to all nine children being obligate homozygotes (**Fig. 4e**). In this family, we challenged homozygotes (n = 6) and non-carriers (n = 7) with a 50 g/m^2^ oral fat load followed by serial blood testing for six hours. *APOC3* p.Arg19Ter carriers had significantly lower post-prandial triglyceride excursions (triglycerides area under the curve 468.3 mg/dL*6 hours vs 1267.7 mg/dL*6 hours; *P* = 1 × 10^−4^) (**Fig. 4f**). These data show that complete lack of APOC3 markedly improves clearance of plasma triglycerides after a fatty meal.

Gene disruption in model organisms followed by phenotypic analysis has been a fruitful approach to understand gene function; here, we extend this concept to the human organism, leveraging naturally-occurring null mutations, consanguinity, and extensive biochemical phenotyping. These results permit several conclusions.

First, power to identify human knockouts is improved with the study of populations with high degrees of consanguinity. Using the observed median inbreeding coefficient of sequenced participants, we estimate that with the sequencing of 200,000 Pakistanis, about 8,754 genes (95% CI, 8,669–8,834) will be completely knocked out in at least one participant (**Fig. 5**). In contrast, with the sequencing of a similar number of outbred individuals of European, East Asian, or African American ancestries, the expected number of genes knocked out is much less at 1,382 (95% CI, 1,339−1,452), 1,423 (95% CI, 1,388−1,457), and 1,822 (95% CI, 1,772−1,871), respectively (minor allele frequency estimates obtained from the Exome Aggregation Consortium, manuscript submitted in parallel). Thus, if a similar number of individuals were sequenced across different ancestries, the number of genes completely knocked out would be nearly six-fold higher in Pakistanis when compared with other outbred populations. For example, among >100,000 participants in Iceland, 1107 homozygous null genes were identified whereas we observe nearly the same number of null genes after analysis of about 7,000 Pakistanis.^3^

**Fig 5.**
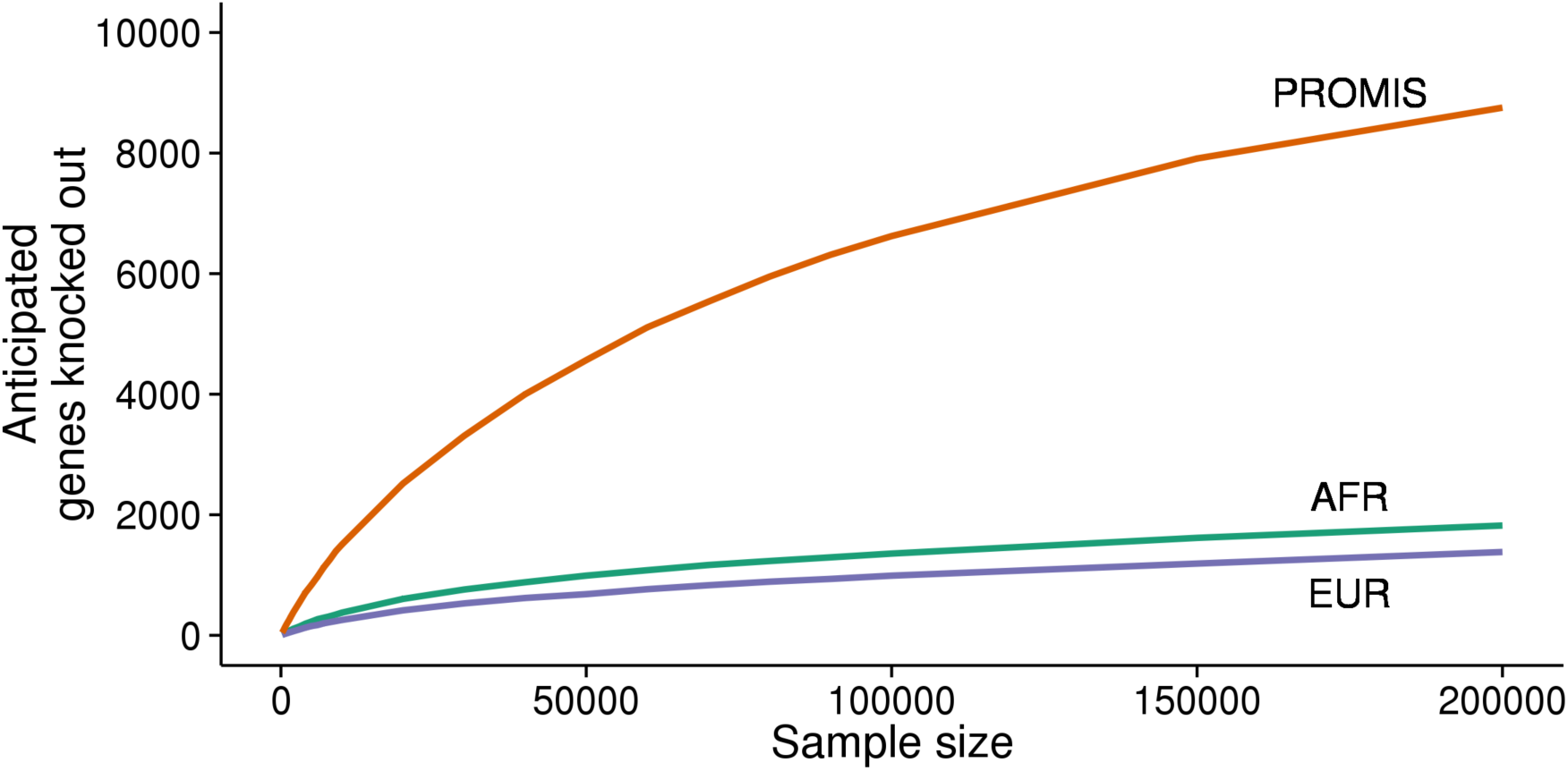
Simulations anticipate many more homozygous null genes in the PROMIS cohort.

Second, dense phenotyping can uncover a range of phenotypic consequences from complete disruption of a gene as observed for *PLA2G7, CYP2F1, A3GALT2* and *NRG4.* Third, recall of complete human knockouts followed by provocative testing may provide physiologic insights. We used this approach to demonstrate that complete lack of apolipoprotein C-III is tolerated and results in both lowered fasting triglyceride concentrations as well as blunted post-prandial lipemia. Finally, to date, most human genetic studies have pursued a phenotype-first (“forward” genetics) approach, beginning with traits of interest followed by genetic mapping. Here, we show that it is now possible to pursue a systematic genotype-first (“reverse” genetics) approach, starting with homozygous null humans followed by methodical examination of a diverse set of traits.

## Methods

### General overview of the Pakistan Risk for Myocardial Infarction Study (PROMIS)

The PROMIS study is designed to investigate determinants of cardiometabolic diseases in Pakistan. Since 2005, the study has enrolled close to 38,000 participants; the present investigation included 7,078 participants. Participants aged 30–80 years were enrolled from nine recruitment centers based in five major urban cities in Pakistan. Type 2 diabetes in the study was defined based on self-report or fasting glucose levels >125 mg/dL or HbA1c > 6.5 % or use of glucose lowering medications. The study was approved by the institutional review board at the Center for Non-Communicable Diseases (IRB: 00007048, I0RG0005843, FWAS00014490) and all participants gave informed consent.

### Phenotype descriptions

Non-fasting blood samples (with the time since last meal recorded) were drawn and centrifuged within 45 minutes of venipuncture. Serum, plasma and whole blood samples were stored at −70°C within 45 minutes of venipuncture. All samples were transported on dry ice to the central laboratory at the Center for Non-Communicable Diseases (CNCD), Pakistan, where serum and plasma samples were aliquoted across 10 different storage vials. Samples were stored at −70°C for any subsequent laboratory analyses. All biochemical assays were conducted in automated auto-analyzers. At CNCD Pakistan, measurements for total-cholesterol, HDL cholesterol, LDL cholesterol, triglycerides, and creatinine were made in serum samples using enzymatic assays; whereas levels of HbA1c were measured using a turbidemetric assay in whole-blood samples (Roche Diagnostics, USA). For further measurements, aliquots of serum and plasma samples were transported on dry ice to the Smilow Research Center, University of Pennsylvania, USA, where following biochemical assays were conducted: apolipoproteins (apoA-I, apoA-II, apoB, apoC-III, apoE) and non-esterified fatty acids were measured through immunoturbidometric assays using kits by Roche Diagnostics or Kamiya; lipoprotein (a) levels were determined through a turbidimetric assay using reagents and calibrators from Denka Seiken (Niigata, Japan); LpPLA2 mass and activity levels were determined using immunoassays manufactured by diaDexus (San Francisco, CA, USA); measurements for insulin, leptin and adiponectin were made using radio-immunoassays by LINCO (MO, USA); levels of adhesion molecules (ICAM-1, VCAM-1, P- and E-Selectin) were determined through enzymatic assays by R&D (Minneapolis, MN, USA); and measurements for C-reactive protein, alanine transaminase, aspartate transaminase, cystatin-C, ferritin, ceruloplasmin, thyroid stimulating hormone, alkaline phosphatase, sodium, potassium, choloride, phosphate, sex-harmone binding globulin were made using enzymatic assays manufactured by Abbott Diagnostics (NJ, USA). Glomerular filtration rate (eGFR) was estimated from serum creatinine levels using the MDRD equation. ApoC-III levels were determined in an autoanalyzer using a commercially available ELISA by Sekisui Diagnostics (Lexington, USA). We also measured the following 52 protein biomarkers by multiplex immunoassay using a customised panel on the Luminex 100/200 instrument by RBM (Myriad Rules Based Medicine, Austin, TX, USA): fatty acid binding protein, granuloctye monocyte colony stimulating factor, granulocyte colony stimulating factor, interferon gamma, interleukin-1 beta, interleukin 1 receptor, interleukin 2, interleukin 3, interleukin 4, interleukin 5, interleukin 6, interleukin 7, interleukin 8, interleukin 10, interleukin 18, interleukin p40, interleukin p70, interleukin 15, interleukin 17, interleukin 23, macrophage inflammatory protein 1 alpha, macrophage inflammatory protein 1 beta, malondialdehyde-modified LDL, matrix metalloproteinase 2, matrix metalloproteinase 3, matrix metalloproteinase 9, nerve growth factor beta, tumor necrosis factor alpha, tumor necrosis factor beta, brain-derived neurotrophic factor, CD40, CD40 ligand, eotaxin, factor VII, insulin-like growth factor 1, lecithin-type oxidized LDL receptor 1, monocyte chemoattractant protein 1, myeloperoxidase, N-terminal prohormone of brain natriuretic peptide, neuronal cell adhesion molecule, pregnancy-associated plasma protein A, soluble receptor for advanced glycation end-products, sortilin, stem cell factor, stromal cell-derived factor 1, thrombomodulin, S100 calcium binding protein B, and vascular endothelial growth factor.

### Laboratory methods for array-based genotyping

As previously described, a genomewide association scan was performed using the Illumina 660 Quad array at the Wellcome Trust Sanger Institute (Hinxton, UK) and using the Illumina HumanOmniExpress at Cambridge Genome Services, UK.^22^ Initial quality control (QC) criteria included removal of participants or single nucleotide polymorphisms (SNPs) that had a missing rate >5%. SNPs with a MAF <1% and a P-value of <10^−7^ for the Hardy-Weinberg equilibrium test were also excluded from the analyses. In PROMIS, further QC included removal of participants with discrepancy between their reported sex and genetic sex determined from the X chromosome. To identify sample duplications, unintentional use of related samples (cryptic relatedness) and sample contamination (individuals who seem to be related to nearly everyone in the sample), identity-by-descent (IBD) analyses were conducted in PLINK.^23^

### Laboratory methods for exome sequencing

Exome sequencing. Exome sequencing was performed at the Broad Institute. Sequencing and exome capture methods have been previously described.^24,25^ A brief description of the methods is provided below.

Receipt/quality control of sample DNA. Samples were shipped to the Biological Samples Platform laboratory at the Broad Institute of MIT and Harvard (Cambridge, MA, USA). DNA concentration was determined by PicoGreen (Invitrogen; Carlsbad, CA, USA) prior to storage in 2D-barcoded 0.75 ml Matrix tubes at −20 ^o^C in the SmaRTStore (RTS, Manchester, UK) automated sample handling system. Initial quality control (QC) on all samples involving sample quantification (PicoGreen), confirmation of high-molecular weight DNA and fingerprint genotyping and gender determination (Illumina iSelect; Illumina; San Diego, CA, USA). Samples were excluded if the total mass, concentration, integrity of DNA or quality of preliminary genotyping data was too low.

**Library construction**. Library construction was performed as previously described^26^, with the following modifications: initial genomic DNA input into shearing was reduced from 3μg to 10–100ng in 50μL of solution. For adapter ligation, Illumina paired end adapters were replaced with palindromic forked adapters, purchased from Integrated DNA Technologies, with unique 8 base molecular barcode sequences included in the adapter sequence to facilitate downstream pooling. With the exception of the palindromic forked adapters, the reagents used for end repair, A-base addition, adapter ligation, and library enrichment PCR were purchased from KAPA Biosciences (Wilmington, MA, USA) in 96-reaction kits. In addition, during the post-enrichment SPRI cleanup, elution volume was reduced to 20 μL to maximize library concentration, and a vortexing step was added to maximize the amount of template eluted.

**In-solution hybrid selection**. 1,973 samples underwent in-solution hybrid selection as previously described^26^, with the following exception: prior to hybridization, two normalized libraries were pooled together, yielding the same total volume and concentration specified in the publication. 5,263 samples underwent hybridization and capture using the relevant components of Illumina’s Rapid Capture Exome Kit and following the manufacturer’s suggested protocol, with the following exceptions: first, all libraries within a library construction plate were pooled prior to hybridization, and second, the Midi plate from Illumina’s Rapid Capture Exome Kit was replaced with a skirted PCR plate to facilitate automation. All hybridization and capture steps were automated on the Agilent Bravo liquid handling system.

**Preparation of libraries for cluster amplification and sequencing**. Following post-capture enrichment, libraries were quantified using quantitative PCR (KAPA Biosystems) with probes specific to the ends of the adapters. This assay was automated using Agilent’s Bravo liquid handling platform. Based on qPCR quantification, libraries were normalized to 2nM and pooled by equal volume using the Hamilton Starlet. Pools were then denatured using 0.1 N NaOH. Finally, denatured samples were diluted into strip tubes using the Hamilton Starlet.

**Cluster amplification and sequencing**. Cluster amplification of denatured templates was performed according to the manufacturer’s protocol (Illumina) using HiSeq v3 cluster chemistry and HiSeq 2000 or 2500 flowcells. Flowcells were sequenced on HiSeq 2000 or 2500 using v3 Sequencing-by-Synthesis chemistry, then analyzed using RTA v.1.12.4.2. Each pool of whole exome libraries was run on paired 76bp runs, with and 8 base index sequencing read was performed to read molecular indices, across the number of lanes needed to meet coverage for all libraries in the pool.

**Read mapping and variant discovery**. Samples were processed from real-time base-calls (RTA v.1.12.4.2 software [Bustard], converted to qseq.txt files, and aligned to a human reference (hg19) using Burrows–Wheeler Aligner (BWA).^27^ Aligned reads duplicating the start position of another read were flagged as duplicates and not analysed. Data was processed using the Genome Analysis ToolKit (GATK v3).^28−30^ Reads were locally realigned around indels and their base qualities were recalibrated. Variant calling was performed on both exomes and flanking 50 base pairs of intronic sequence across all samples using the HaplotypeCaller (HC) tool from the GATK to generate a gVCF. Joint genotyping was subsequently performed and ‘raw’ variant data for each sample was formatted (variant call format (VCF)). SNVs and indel sites were initially filtered after variant calibration marked sites of low quality that were likely false positives.

**Data analysis QC**. Fingerprint concordance between sequence data and fingerprint genotypes was evaluated. Variant calls were evaluated on both bulk and per-sample properties: novel and known variant counts, transition–transversion (TS–TV) ratio, heterozygous–homozygous non-reference ratio, and deletion/insertion ratio. Both bulk and sample metrics were compared to historical values for exome sequencing projects at the Broad Institute. No significant deviation of from historical values was noted.

### Data processing and quality control of exome sequencing

**Variant annotation**. Variants were annotated using Variant Effect Predictor^31^ and the LOFTEE^8^ plugin to identify protein-truncating variants predicted to disrupt the respective gene’s function with “high confidence.” Each allele at polyallelic sites was separately annotated.

**Sample level quality control**. We performed quality control of samples using the following steps. For quality control of samples, we used bi-allelic SNVs that passed the GATK VQSR filter and were on genomic regions targeted by both ICE and Agilent exome captures. We removed samples with discordance rate > 10% between genotypes from exome sequencing with genotypes from array-based genotyping and samples with sex mismatch between inbreeding coefficient on chromosome X and fingerprinting. We tested for sample contamination using the verifyBamID software, which examines the proportion of non-reference bases at reference sites, and excluded samples with high estimated contamination (FREEMIX scores > 0.2).^32^ After removing monozygotic twins or duplicate samples using the KING software^33^, we removed outlier samples with too many or too few SNVs (>17,500 total variants for Agilent-captured samples or >18,000 for ICE-captured samples; <12,000 total variants; >600 singletons; and >400 doubletons). We removed those with extreme overall transition-to-transversion ratios (>4 or <3) and heterozygosity (heterozygote-to-homozygote ratio >6 or <2). Finally, we removed samples with high missingness (>0.05).

**Variant level quality control**. Variant score quality recalibration was performed separately for SNVs and indels use the GATK VariantRecalibrator and ApplyRecalibration to filter out variants with lower accuracy scores. To further reduce the rate of inaccurate variant calls, we further filtered out SNVs with low average quality (quality per depth of coverage (QD) < 2) and a high degree of missingness (> 20 %), and indels also with low average quality (quality per depth of coverage (QD) < 3) and a high degree of missingness (> 20 %).

### Methods for inbreeding analyses

**Array-derived runs of homozygosity**. Analyses were conducted in PLINK^23^ using genome-wide association (GWAS) data in PROMIS and HapMap 3 populations. Segments of the genome that were at-least 1.5 Mb long, had a SNP density of 1 SNP per 20 kb and had 25 consecutive homozygous SNPs (1 heterozygous and/or 5 missing SNPs were permitted within a segment) were defined to be in a homozygous state (or referred as “runs of homozygosity” (ROH)), as described previously.^34^ Homozygosity was expressed as the percentage of the autosomal genome found in a homozygous state, and was calculated by dividing the sum of ROH length within each individual by the total length of the autosome in PROMIS and HapMap 3 populations respectively. To investigate variability in homozygosity explained by parental consanguinity, the difference in R^2^ is reported for a linear regression model of homozygosity including and excluding parental consanguinity on top of age, sex and the first 10 principal components derived from the typed autosomal GWAS data.

**Sequencing-derived coefficient of inbreeding**. We compared the coefficient of inbreeding distributions of 7,078 exome sequenced PROMIS participants with 15,248 participants (European ancestry = 12,849, and African ancestry = 2,399) who were exome sequenced at the Broad Institute (Cambridge, MA) from the Myocardial Infarction Genetics consortium.^25^ We extracted approximately 5,000 high-quality polymorphic SNVs in linkage equilibrium present on both target intervals that passed variant quality control metrics based on HapMap 3 data.^35^ Using PLINK, we estimated the coefficient of inbreeding separately within each ethnicity group.^23^ The coefficient of inbreeding was estimated as the observed degree of homozygosity compared with the anticipated homozygosity derived from an estimated common ancestor.^36^ The Wilcoxon-Mann-Whitney test was used to test whether PROMIS participants had different median coefficients of inbreeding compared to other similarly sequenced outbred individuals and whether the median coefficient of inbreeding was different between PROMIS participants who reported parental relatedness versus not. A two-sided *P* of 0.05 was the pre-specified threshold for statistical significance.

### Methods for sequencing projection analysis

To compare the burden of unique completely inactivated genes in the PROMIS cohort with outbred cohorts of diverse ethnicities, we extracted the minor allele frequencies (maf) of “high confidence” loss-of-function mutations observed in PROMIS, and in European, African, and East Asian ancestry participants from the Exome Aggregation Consortium (ExAC r0.3; exac.broadinstitute.org). For each gene and for each ethnicity, the combined minor allele frequency (cmaf) of rare (maf < 0.1%) “high confidence” loss-of-function mutations was calculated. We then simulated the number of unique completely inactivated genes across a range of sample sizes per ethnicity and PROMIS. The expected probability of observing complete inactivation (two null copies in an individual) of a gene was calculated as (1 − *F*) * *cmaf* ^2^ + *F * cmaf,* which accounts for allozygous and autozygous, respectively, mechanisms for complete genie knockout. F, the inbreeding coefficient, is defined as *F =* 1 − (*expected heterozygosity rate / observed heterozygosity rate*). For PROMIS, the median F inbreeding coefficient (0.016) was used for estimation. Down-sampling within the observed sample size for both high-confidence null mutations and synonymous variants did not deviate significantly from the expected trajectory. For a range of sample sizes (0–200,000), each gene was randomly sampled under a binomial distribution (*X ~ B*(*n, cmaf*)) and it was determined if the gene was successfully sampled at least once. To refine the estimated count of unique genes per sample size, each sampling was replicated ten times.

### Methods for constraint score analysis

We sought to determine whether the observed homozygous null genes were under less evolutionary constraint by first obtaining constraint loss of function constraint scores derived from the Exome Aggregation Consortium (Lek M et al, in preparation).^9,10^ Briefly, we used the number of observed and expected rare (MAF < 0.1%) loss of function variants per gene to determine to which of three classes it was likely to belong: null (observed variation matches expectation), recessive (observed variation is ~50% expectation), or haploinsufficient (observed variation is <10% of expectation). The probability of being loss of function intolerant (pLI) of each transcript was defined as the probability of that transcript falling into the haploinsufficient category. Transcripts with a pLI ≥ 0.9 are considered very likely to be loss of function intolerant; those with pLI ≤ 0.1 are not likely to be loss of function intolerant. A list of 961 genes were randomly sampled from a list of sequenced genes 1,000 times and the proportion of loss of function intolerant genes compared to the proportion of the observed homozygous null genes was compared using the chi square test. The likelihood that the distribution of the test statistics deviated from the null was ascertained.

### Methods for rare variant association analysis

**Recessive model association discovery**. We sought to determine whether complete loss-of-function of a gene was associated with a dense array of phenotypes. We extracted a list of individuals per gene who were homozygous for a high confidence null allele that was rare (minor allele frequency < 1 %) in the cohort. From a list of 961 genes where there was at least one participant homozygous null and a list of 201 traits, we initially considered 192,960 gene-trait pairings. To reduce the likelihood of false positives, we only considered gene-trait pairs where there were at least two homozygous nulls per gene phenotyped for a given trait yielding 15,263 gene-trait pairs for analysis.

For all analyses, we constructed generalized linear models to test whether complete loss of function versus non-carriers was associated with trait variation. A logit link was used for binomial outcomes. Right-skewed continuous traits were natural log transformed. Age, sex, and myocardial infarction status were used as covariates in all analyses. We extracted principal components of ancestry using EIGENSTRAT to control for population stratification in all analyses.^37^ For lipoprotein-related traits, the use of lipid-lowering therapy was used as a covariate. For glycemic biomarkers, only non-diabetics were used in the analysis. The *P* threshold for statistical significance was 0.05 / 15,263 = 3 × 10^−6^.

**Heterozygote association replication**. We hypothesized that some of the associations for homozygous nulls will display a more modest effect for heterozygous nulls. Thus, the aforementioned analyses were performed comparing heterozygous nulls to non-carriers for the fourteen homozygous null-trait associations that surpassed prespecified statistical significance. A *P* of 0.05 / 13 = 0.004 was set for statistical significance for these restricted analyses.

**Association for single genic homozygotes**. We performed an exploratory analysis of gene-trait pairs where there was only one phenotyped homozygous null. We performed the above association analyses for genes where there was only one homozygous null phenotyped for a given trait and we focused on those with the most extreme standard Z score statistics (|Z score| > 5) from the primary association analysis and required that there to also be nominal evidence for association (*P* < 0.05) in heterozygotes as well to maximize confidence in an observed single homozygous null-trait association.

### Methods for recruitment and phenotyping of an *APOC3* p.Arg19Ter proband and relatives

**Methods for Sanger sequencing**. We collected blood samples from a total of 28 subjects, including one of the four *APOC3* p.Arg19Ter homozygous participants along with 27 of his family and community members for DNA extraction and separated into plasma for lipid and apolipoprotein measurements. All subjects were consented prior to initiation of the studies (IRB: 00007048 at the Center for Non-Communicable Diseases, Paksitan). DNA was isolated from whole blood using a reference phenol-chloroform protocol.^38^ Genotypes for the p.Arg19Ter variant were determined in all 28 participants by Sanger sequencing. A 685 bp region of the *APOC3* gene including the base position for this variant was amplified by PCR (Expand HF PCR Kit, Roche) using the following primer sequences: Forward primer CTCCTTCTGGCAGACCCAGCTAAGG, Reverse primer CCTAGGACTGCTCCGGGGAGAAAG. PCR products were purified with Exo-SAP-IT (Affymetrix) and sequenced via Sanger sequencing using the same primers.

**Oral fat tolerance test**. Six non-carriers and seven homozygotes also participated in an oral fat tolerance test. Participants fasted overnight and then blood was drawn for measurement of baseline fasted lipids. Following this, participants were administered an oral load of heavy cream (50 g fat per square meter of body surface area as calculated by the method of Mosteller^39^). Participants consumed this oral load within a time span of 20 minutes and afterwards consumed 200 mL of water. Blood was drawn at 2, 4, and 6 hours after oral fat consumption as done previously.^40,41^ All lipid and apolipoprotein measurements from these plasma samples were determined by immunoturbidimetric assays on an ACE Axcel Chemistry analyzer (Alfa Wasserman). A comparisons of area-under-the curve triglycerides was performed between *APOC3* p.Arg19Ter homozygotes and non-carriers using a two independent sample Student’s t test; *P* < 0.05 was considered statistically significant.

**Figure Legends**

**Fig 1**. Consanguinity leads to regions of genomic segments that are identical by descent and can be observed as runs of homozygosity. Using genome-wide array data in 17,744 PROMIS participants and reference samples from the International HapMap 3, the burden of runs of homozygosity (minimum 1.5 Mb) per individual was derived and population-specific distributions are displayed, with outliers removed. This highlights the higher median runs of homozygosity burden in PROMIS and the higher proportion of individuals with very high burdens.

**Fig 2**. The site-frequency spectrum of synonymous, missense, and high-confidence null mutations is represented. Points represent the proportion of variants within a 1 × 10^−4^ minor allele frequency bin for each variant category. Lines represent the cumulative proportions of variants categories. The bottom inset highlights that most null variants are often seen in no more than one or two individuals. The top inset highlights that virtually all null mutations are very rare.

**Fig 3**. **a**, The distribution of F inbreeding coefficient of PROMIS participants is compared to those of outbred samples of African (AFR) and European (EUR) ancestry. **b**, The burden of homozygous null genes per individual is correlated with coefficient of inbreeding.

**Fig 4**. **a.-d.** Among all sequenced participants, apolipoprotein C-III, triglycerides, HDL cholesterol and LDL cholesterol distributions are displayed by *APOC3* null genotype status. Apolipoprotein C-III concentration is displayed on a logarithmic base 10 scale. **e**. One of the four *APOC3* null homozygotes and several family members were recruited for genotyping *APOC3* p.Arg19Ter. The proband was married to another null homozygote and has nine obligate homozygote children. Given the extensive first-degree unions, the pedigree is simplified for clarity. **f**. *APOC3* p.Arg19Ter homozygotes and non-carriers within the recruited family participated in an oral 50 g/m^2^ fat tolerance test. Homozygotes had decreased baseline and post-prandial triglyceride concentrations.

**Fig 5**. The anticipated number of unique homozygous null genes observed with increasing sample sizes sequenced in PROMIS compared with similar African (AFR) and European (EUR) sample sizes using observed allele frequencies and degree of inbreeding.

**Supplementary Information** is linked to the online version of the paper at www.nature.com/nature.

## Acknowledgements

Dr. Saleheen is supported by grants from the National Institutes of Health, the Fogarty International, the Wellcome Trust, the British Heart Foundation, and Pfizer. Dr. Natarajan is supported by the John S. LaDue Memorial Fellowship in Cardiology from Harvard Medical School. Dr. Kathiresan is supported by grants from the National Institutes of Health (R01HL107816), the Donovan Family Foundation, and Fondation Leducq. Exome sequencing was supported by a grant from the NHGRI (5U54HG003067–11) to Drs. Gabriel and Lander. Dr. MacArthur is supported by a grant from the National Institutes of Health (R01GM104371). In recognition for PROMIS fieldwork and support, we also acknowledge contributions made by the following: Mohammad Zeeshan Ozair, Usman Ahmed, Abdul Hakeem, Hamza Khalid, Kamran Shahid, Fahad Shuja, Ali Kazmi, Mustafa Qadir Hameed, Naeem Khan, Sadiq Khan, Ayaz Ali, Madad Ali, Saeed Ahmed, Muhammad Waqar Khan, Muhammad Razaq Khan, Abdul Ghafoor, Mir Alam, Riazuddin, Muhammad Irshad Javed, Abdul Ghaffar, Tanveer Baig Mirza, Muhammad Shahid, Jabir Furqan, Muhammad Iqbal Abbasi, Tanveer Abbas, Rana Zulfiqar, Muhammad Wajid, Irfan Ali, Muhammad Ikhlaq, Danish Sheikh, Muhammad Imran, Matthew Walker, Nadeem Sarwar, Sarah Venorman, Robin Young, Adam Butterworth, Hannah Lombardi, Binder Kaur and Nasir Sheikh. Fieldwork in the PROMIS study has been supported through funds available to investigators at the Center for Non-Communicable Diseases, Pakistan and the University of Cambridge, UK.

## Author Contributions

Sample recruitment and phenotyping was performed by D.S., P.F., J.D., A.R., M.Z., M.S., M.F., A.I., N.K.S., S.A., F.M., M.I., S.A., K.T., N.H.M., K.S.Z., N.Q., M.I., S.Z.R., F.M., K.M., and N.A., D.S., P.F., J.D., and W.Z. performed array-based genotyping and runs-of-homozygosity analyses. Exome sequencing was coordinated by D.S., N.G., S.G., E.S.L., D.J.R., and S.K., P.N., W.Z., H.H.W., and R.D. performed exome sequencing quality control and association analyses. P.N., K.J.K., A.H.O., and D.G.M. performed variant annotation. D.S., S.K. and D.J.R. performed confirmatory genotyping and lipoprotein biomarker assays. D.S. and A.R. conducted recall based studies for the APOC3 knockouts. P.N. and M.J.D. performed bioinformatics simulations. P.N. and K.E.S. performed constraint score analyses. D.S., P.N., and S.K. designed the study and wrote the paper. D.S. and P.N. contributed equally. All authors discussed the results and commented on the manuscript.

**Author Information** Summaries of all null variants observed in a homozygous are in the online Supplement. They are additionally, with all observed protein-coding variation, publicly available in the Exome Aggregation Consortium browser (exac.broadinstitute.org). Reprints and permissions information is available at www.nature.com/reprints. The authors do not declare competing financial interests. Correspondence and requests for materials should be addressed to D.S. or S.K.

